# MEYE: Web-app for translational and real-time pupillometry

**DOI:** 10.1101/2021.03.09.434438

**Authors:** Raffaele Mazziotti, Fabio Carrara, Aurelia Viglione, Leonardo Lupori, Luca Lo Verde, Alessandro Benedetto, Giulia Ricci, Giulia Sagona, Giuseppe Amato, Tommaso Pizzorusso

## Abstract

Pupil dynamics alterations have been found in patients affected by a variety of neuropsychiatric conditions, including autism. Studies in mouse models have used pupillometry for phenotypic assessment and as a proxy for arousal. Both in mice and humans, pupillometry is non-invasive and allows for longitudinal experiments supporting temporal specificity, however its measure requires dedicated setups. Here, we introduce a Convolutional Neural Network that performs on-line pupillometry in both mice and humans in a web app format. This solution dramatically simplifies the usage of the tool for non-specialist and non-technical operators. Because a modern web browser is the only software requirement, this choice is of great interest given its easy deployment and set-up time reduction. The tested model performances indicate that the tool is sensitive enough to detect both spontaneous and evoked pupillary changes, and its output is comparable with state-of-the-art commercial devices.

## Introduction

Pupillometry, the measurement of pupil size fluctuations over time, provides useful insights in clinical settings and basic research activity. Light level is the primary determinant of pupil size, even though non-light-driven pupil fluctuations, widely assumed as an indicator of arousal through Locus Coeruleus (LC) activity, can be used to index brain state across species [1–3]. Higher cognitive and emotional processes are also able to evoke tonic or phasic pupillary changes, such as attention [4], memory load [5], novelty [6–8], pain [9–11], and more general cortical sensory processing [2,12] in humans and in animal models.

A growing body of work shows how pupillometry can be used as a possible biomarker for numerous neurological and psychiatric conditions in early development and adult subjects [13– 27]. Spontaneous and voluntary modulation of pupil fluctuations have also been used to facilitate Human-Computer Interaction in normal subjects [28–30] and patients with severe motor disabilities. For example, pupil dynamics is used to assess communication capability in Locked-in Syndrome, a crucial factor for the determination of a minimally conscious state [31,32]. Pupillometry is also becoming a valuable tool for child neurology, to facilitate risk assessment in infants. For example, the Pupil Light Reflex (PLR) during infancy seems to predict the later diagnosis and severity of Autism Spectrum Disorders (ASD) [27]. Intriguingly, pupil alterations are also present in several ASD mouse models [26].

Pupillometry has several advantages as compared with other physiological methods: it is non-invasive and can be performed by non-specialized personnel on non-collaborative and preverbal subjects (like infants), allowing the design of longitudinal experiments to permit temporal specificity. More importantly, it can be conducted similarly across different species from mice to humans, guaranteeing maximal translatability of the protocols and results [13,16,26]. Given these assumptions, it is vital to introduce a simple, versatile tool used in a range of settings, from the laboratory to the clinical or even domestic environment. Available open source methods require complicated steps for the installation and configuration of custom software not suitable for non-technical operators. Moreover, these tools were tested exclusively in one species (mice [33], humans [34]), and none of them were applied in cognitive experiments that usually involve small pupil changes associated with high variability.

In this work, we have developed a deep learning tool called MEYE, using convolutional neural networks (CNNs) to detect and measure real-time changes in pupil size both in humans and mice in different experimental conditions. Furthermore, MEYE web app, performs pupil area quantification and blink detection, all within a single network. By embedding artificial intelligence algorithms in a web browser to process real-time webcam streams or videos of the eye, MEYE can be used by non-technical operators, opening the possibility to perform pupillometry widely, cost-effectively, and in a high-throughput manner. This architecture is resistant to different illumination conditions, allowing the design of basic neuroscience experiments in various experimental settings, such as behavior coupled with electrophysiology or imaging like 2-photon microscopy. To describe the performances of MEYE web app in different settings, we tested the app in both mice and humans. In mice we recorded both running speed and pupil size during auditory stimulation. In humans we tested MEYE capabilities to detect the PLR. Furthermore, we performed a visual oddball paradigm [35–37], comparing pupil size and eye position measurements obtained from MEYE with one of the most used commercial eye-tracker systems: the EyeLink 1000. Finally we released a dataset of more than 11897 eye images that can be used to train other artificial intelligence tools.

## Methods

### Datasets

For this study, we collected a dataset (Fig. 1 A) composed of 11897 grayscale images of humans (4285) and mouse (7612) eyes. The pictures’ majority depicts mouse eyes during head-fixation sessions (HF: 5061) in a dark environment using infrared (IR, 850 nm) light sources. In this environment, the pupil is darker than the rest of the image. We also collected mouse eyes (2P: 2551) during 2-photon Ca2+ imaging. In this particular condition, the pupil is inverted in color and tends to be brighter than the iris. Finally, we acquired images of human eyes in IR light (H: 4285) during virtual reality experiments (wearing a headset for virtual reality), using an endoscopic camera (www.misumi.com.tw/). The dataset contains 1596 eye blinks, 841 images in the mouse, and 755 photos in the human datasets. Five human raters segmented the pupil in all pictures (one per image), using custom labeling scripts implemented in Matlab or Python, by manual placement of an ellipse or polygon over the pupil area. Raters flagged blinks using the same code.

**Fig. 1:**
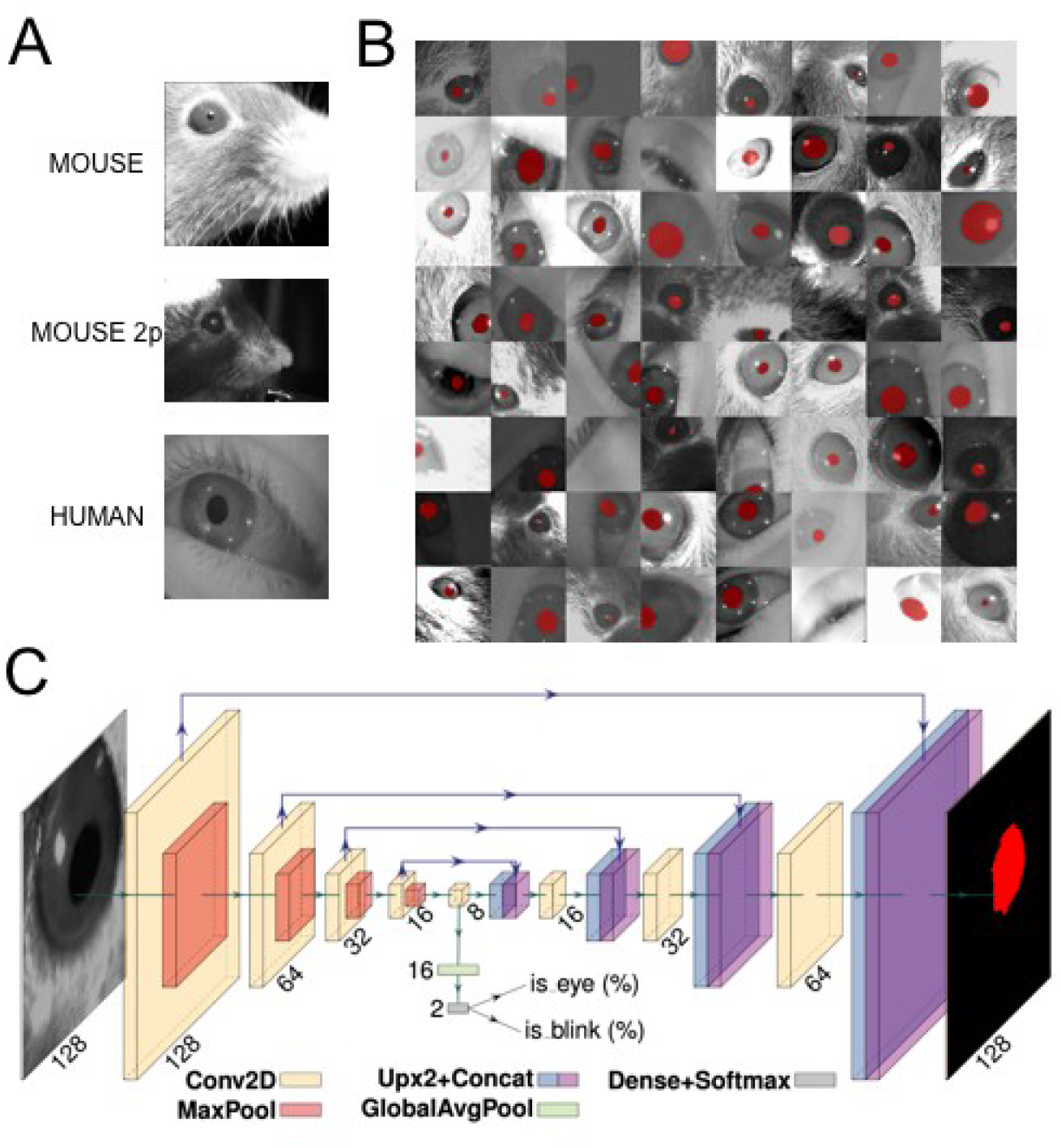
Dataset, CNN architecture, and performances. **A:** examples of images taken from the dataset. The first image depicts a head-fixed mouse with dark pupils, the second one is a head-fixed mouse with a bright pupil, during 2-photon microscope sessions. The last image is a human eye taken during experiments wearing virtual reality goggles. **B:** 64 examples of data augmentation fed to CNN. The images are randomly rotated, cropped, flipped (horizontally or vertically), and changed in brightness/contrast/sharpness. **C:** CNN architecture with an encoder-decoder “hourglass” shape. The encoder part comprises a sequence of convolutional layers. Starting from the last encoder output, the decoder part iteratively upsamples and fuses feature maps with corresponding encoder’s maps, to produce the output pixel map. The pixel probability map and eye/blink probabilities are computed by applying the sigmoid activation to the network outputs element-wise.

**Fig. 2:**
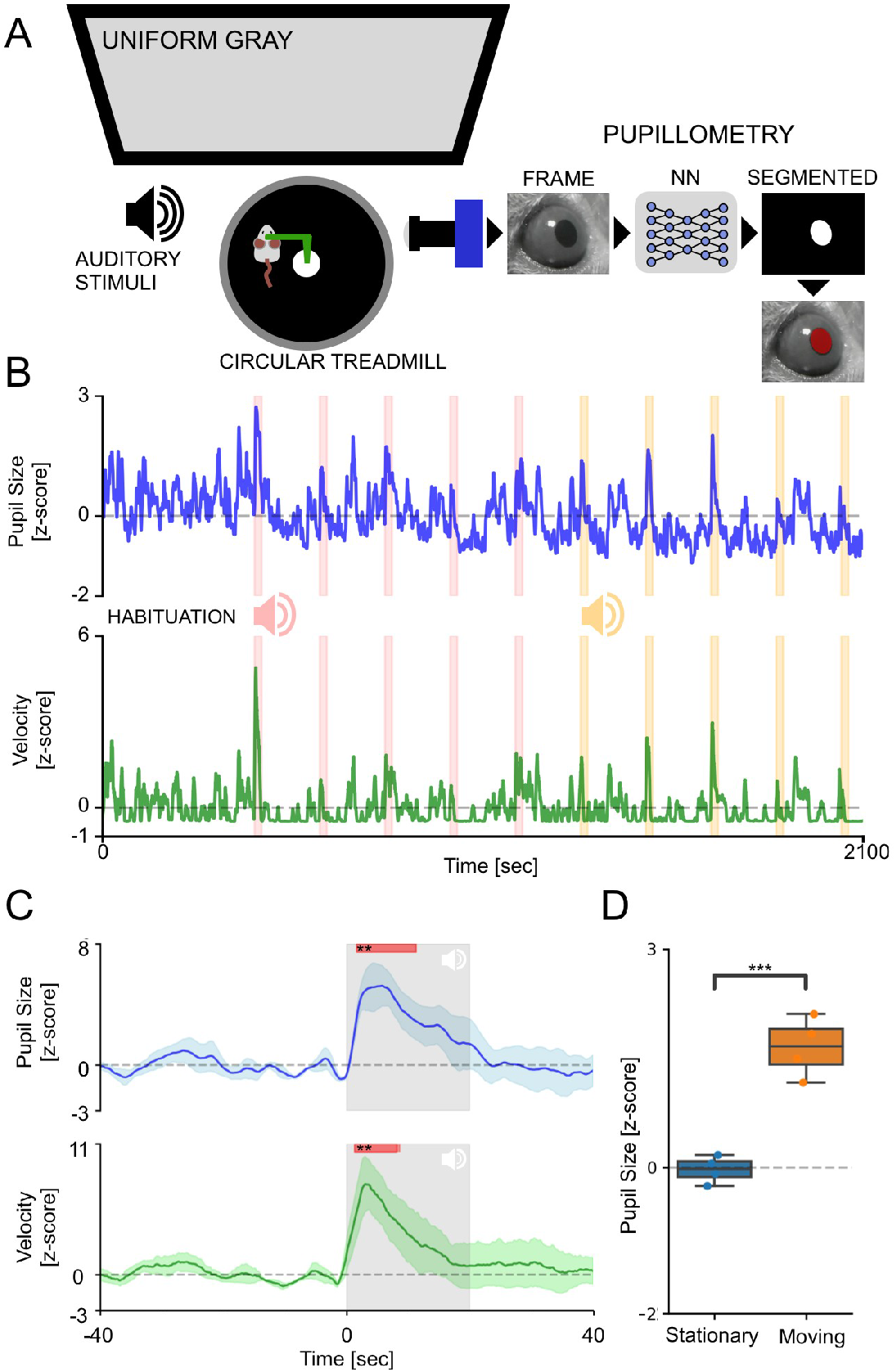
Pupillometry in head-fixed mice. **A:** Setup for head-fixed pupillometry in the awake mouse. The mouse is head-fixed to a custom made metal arm equipped with a 3D printed circular treadmill to monitor running behavior. In the meantime, pupillometry is performed using CNN. **B:** The average fluctuation of pupillometry and velocity in all experimental mice. Dashed pink and yellow areas represent the onset and duration of auditory stimuli. Evoked peaks in both pupil size (blue line) and velocity (green line) are clearly noticeable during auditory stimulation. **C:** Average event-related transients for both pupil size and velocity. Gray areas represent stimulus onset and duration. Red areas represent statistically significant data points with respect to random permutation testing. **D:** Sensibility of the system to detect spontaneous arousal fluctuations. Average pupil size is significantly affected by the behavioral states of the animal. During running epochs (Moving) the pupil is significantly more dilated than during the resting state (Stationary).

### CNN Architecture

The adopted CNN (Fig. 1 C) takes a grayscale 128×128 image as input and produces three outputs: a) a 128×128 probability map of each pixel belonging to the pupil, b) the probability the image contains an eye, and c) the probability the image depicts a blinking eye. We adopted a U-Net variant [38], a widely used CNN in image segmentation tasks. The model has an encoder-decoder “hourglass” architecture; the encoder part comprises a sequence of convolutional layers with ReLU activation and 2×2 max pooling operation, each halving the spatial resolution of feature maps at every layer; this produces a sequence of feature maps of diminishing spatial dimensions that provides both spatially local information and global context for the subsequent steps. Starting from the last encoder output, the decoder part iteratively upsamples and fuses feature maps with corresponding encoder maps, using convolutional layers, to produce the output pixel map. All convolutional layers have 16 3×3 kernels and pad their input to obtain a same-shaped output. Upsampling and downsampling operations have factor 2. Eye and blink probabilities are predicted by an additional branch that applies global average pooling and a two-output fully-connected layer to the bottleneck feature map. The pixel probability map and eye/blink probabilities are computed by applying the sigmoid activation to the network outputs element-wise.

### Augmentation, Training, and Validation

We randomly split the dataset into training, validation, and test subsets following a 70/20/10% split. We perform strong data augmentation during the training phase by applying random rotation, random cropping, random horizontal and vertical flipping, and random brightness/contrast/sharpness changes; images are resized to 128×128 before feeding them to the CNN (Fig. 1 B).

For validation and test images, we take a 128×128 crop centered on the pupil. We compute the binary cross-entropy for all outputs (pixels and eye/blink logits) and take the sum as the loss function to minimize. The network is trained with the AdaBelief optimizer [39] for 750 epochs with a learning rate of 0.001. The best performing snapshot on the validation set is selected and evaluated on the test set.

### MEYE: Web-browser Tool

We built a web app for pupillometry on recorded or live-captured videos harnessing our model as the core component. The trained model has been converted to a web-friendly format using *TensorFlow*.*js*, thus enabling predictions on the user machine using a web browser.

This choice greatly facilitates the deployment and reduces set-up time, as a modern web browser is the only minimum requirement. Once loaded, an internet connection is not mandatory, as no data leaves the user’s browser, and all the processing is performed on the user’s machine. This implies that performance greatly depends on the user’s hardware; if available, hardware (GPU) acceleration is exploited automatically by *TensorFlow*.*js*. In our tests, a modern laptop shipping an Intel(R) Core(TM) i7-9750H 2.60GHz CPU and an Intel(R) UHD Graphics 630 GPU can process up to 28 frames per second.

The web app also offers additional features that facilitate the recording process, such as:

- Processing of pre-recorded videos or real-time video streams captured via webcam;
- ROI placement via user-friendly web UI (drag&drop) and automatic repositioning following tracked pupil center;
- Embedded tunable post-processing (map thresholding and refinement via mathematical morphology);
- Support for registering trigger events;
- Live plotting of pupil area and blink probability;
- Data export in CSV format including: pupil area, blink probability, eye position and trigger channels.

### Behavioral Experiments on Mice

#### Animal Handling

Mice were housed in a controlled environment at 22 C with a standard 12-h light-dark cycle. During the light phase, a constant illumination below 40 lux from fluorescent lamps was maintained. Food (standard diet, 4RF25 GLP Certificate, Mucedola) and water were available ad libitum and changed weekly. Open-top cages (36.5×20.7×14 cm; 26.7×20.7×14 cm for up to 5 adult mice or 42.5×26.6×15.5 cm for up to 8 adult mice) with wooden dust-free bedding were used. All the experiments were carried out following the directives of the European Community Council and approved by the Italian Ministry of Health (1225/2020-PR). All necessary efforts were made to minimize both stress and the number of animals used. The subjects used in this work were three female C57BL/6J mice at 3 months of age.

#### Surgery

The mouse was deeply anesthetized using isoflurane (3% induction, 1.5% maintenance). Then it was mounted on a stereotaxic frame through the use of ear bars. Prilocaine was used as a local anesthetic for the acoustic meatus. The eyes were treated with a dexamethasone-based ophthalmic ointment (Tobradex, Alcon Novartis) to prevent cataract formation and keep the cornea moist. Body temperature was maintained at 37 degrees using a heating pad monitored by a rectal probe. Respiration rate and response to toe pinch were checked periodically to maintain an optimal level of anesthesia. Subcutaneous injection of Lidocaine (2%) was performed prior to scalp removal. Skull surface was carefully cleaned and dried, and a thin layer of cyanoacrylate is poured over the exposed skull to attach a custom made head post that is composed of a 3D printed base equipped with a glued set screw (12 mm long, M4 thread, Thorlabs: SS4MS12). The implant is secured to the skull using cyanoacrylate and UV curing dental cement (Fill Dent, Bludental). At the end of the surgical procedure, the mice recovered in a heated cage. After 1 hour, mice were returned to their home cage. Paracetamol was used in the water as antalgic therapy for three days. We wait seven days before performing head-fixed pupillometry to provide sufficient time for the animal to recover.

#### Head Fixation

In the awake mouse head-fixation experiments, we employed a modified version of the apparatus proposed by Silasi et al. [40], equipped with a 3D printed circular treadmill (diameter: 18cm). Components listed in Table1. A locking ball socket mount (TRB1/M) is secured to an aluminum breadboard (MB2020/M) using two optical posts (TR150/M-P5) and a right angle clamp (RA90/M-P5). The circular treadmill is blocked between the base plate pillar rod and the optical post through a ball-bearing element (BU4041, BESYZY) to allow the disk’s spinning with low effort. To couple the head-fixing thread on the mouse to the locking ball, an ER025 post was modified by re-tapping one end of it with M4 threads to fit the ball and socket mount. Velocity is detected using an optical mouse under the circular treadmill. Pupillometry is performed using a USB camera (oCam-5CRO-U, Withrobot) equipped with a 25 mm M12 lens connected to a Jetson *AGX Xavier Developer Kit* (NVIDIA) running a custom Python3 script (30fps). The Jetson hardware is connected with an Arduino UNO through GPIO digital connection. The Arduino UNO manages the auditory stimuli through a speaker (W3-1364SA 3”, Tang Band).

#### Behavioral Procedure

Mice were handled for 5 minutes each day during the week preceding the experiments; then, they were introduced gradually to head-fixation for an increasing amount of time for five days. During Day 1 and 2, we performed two sessions of 10 minutes of head-fixation, one in the morning and one in the afternoon. On Day 3, we performed one session of twenty minutes, Day 4 thirty minutes, and Day 5 thirty-five minutes. Each recording started with 5 minutes of habituation. We exposed the animal to the auditory stimuli during the last day. During each head-fixation session, a curved monitor (24 inches Samsung, CF390) is placed in front of the animal showing a uniform gray with a mean luminance of 8.5 cd/m2. The frequency of tone 1 is 3000Hz, tone 2 is 4000Hz, both at 70dB, 10 seconds duration, and 120 seconds of interstimulus.

#### Data Analysis

Data has been analyzed using Python 3 and Jupyter notebooks. Correlation has been performed using *pingouin*.*corr* (spearman method). Permutation tests were carried out permuting single subject samples for 3000 times using the function *scipy*.*random*.*permutation* to calculate mean and standard deviation of the chance level. Then each sample of the ERT was compared with the corresponding null hypothesis distribution using *scipy*.*stats*.*norm*.*cdf*. All the obtained p-values were corrected for multiple comparisons using the Benjamini/Hochberg FDR correction of *pingouin*.*multcomp* Python function. T-tests were performed using the *pingouin*.*ttest* Python function. Eyes Movements comparison is carried out normalizing (in the range between -1 and 1) data from both setups, upsamplig MEYE data from 15 to 1000 fps using linear interpolation and then calculating the Mean Absolute Error (MAE), performed using the Python function *sklearn*.*metrics*.*mean_absolute_error*.

### Behavioral Experiments on Humans

#### PLR

Pupillometry has been performed using a MacBook Pro (Retina, 13-inch, Early 2015, Intel Core i5 Dual-core 2.7GHz, 8GB of RAM, Intel Iris Graphics 6100 1536 MB) running MEYE application on Firefox (84.0). The tool is able to compute online pupil size quantification, plotting the instantaneous pupil area and saving the results on file. Furthermore the tool accepts four independent manual push button triggers (keys T or Y on the keyboard). This feature allowed us to annotate stimulation events. A USB IR webcam (Walfront5k3psmv97x, Walfront) equipped with a Varifocal 6-22mm M12 objective (149129, Sodial) was used to acquire images of the eye. The camera is equipped with 6 IR LEDs to illuminate the eye uniformly, optimising contrast between the iris and the pupil. Photic stimulation is delivered using an Arduino Due (Arduino) microcontroller connected via USB to the notebook and programmed to emulate a keyboard. The Arduino emulates a keyboard (using the *keyboard*.*h* library) to send event triggers to MEYE in the form of keystroke events. The microcontroller drives a stripe of four LEDs (WS2813, WorldSemi) using the *FastLED*.*h* library, flashing bright white light for 500 ms with an interstimulus of 5 seconds (Fig. 3 A). The Subject sat in front of a monitor screen (24 inches Samsung, CF390) at a distance of 60 cm, with the head stabilized by a chin rest and instructed to maintain fixation on a small dot presented in the center of the screen for the whole duration of the recording (57 seconds). A total of 10 flash stimuli have been presented through the strip of LED mounted above the screen.

**Fig. 3:**
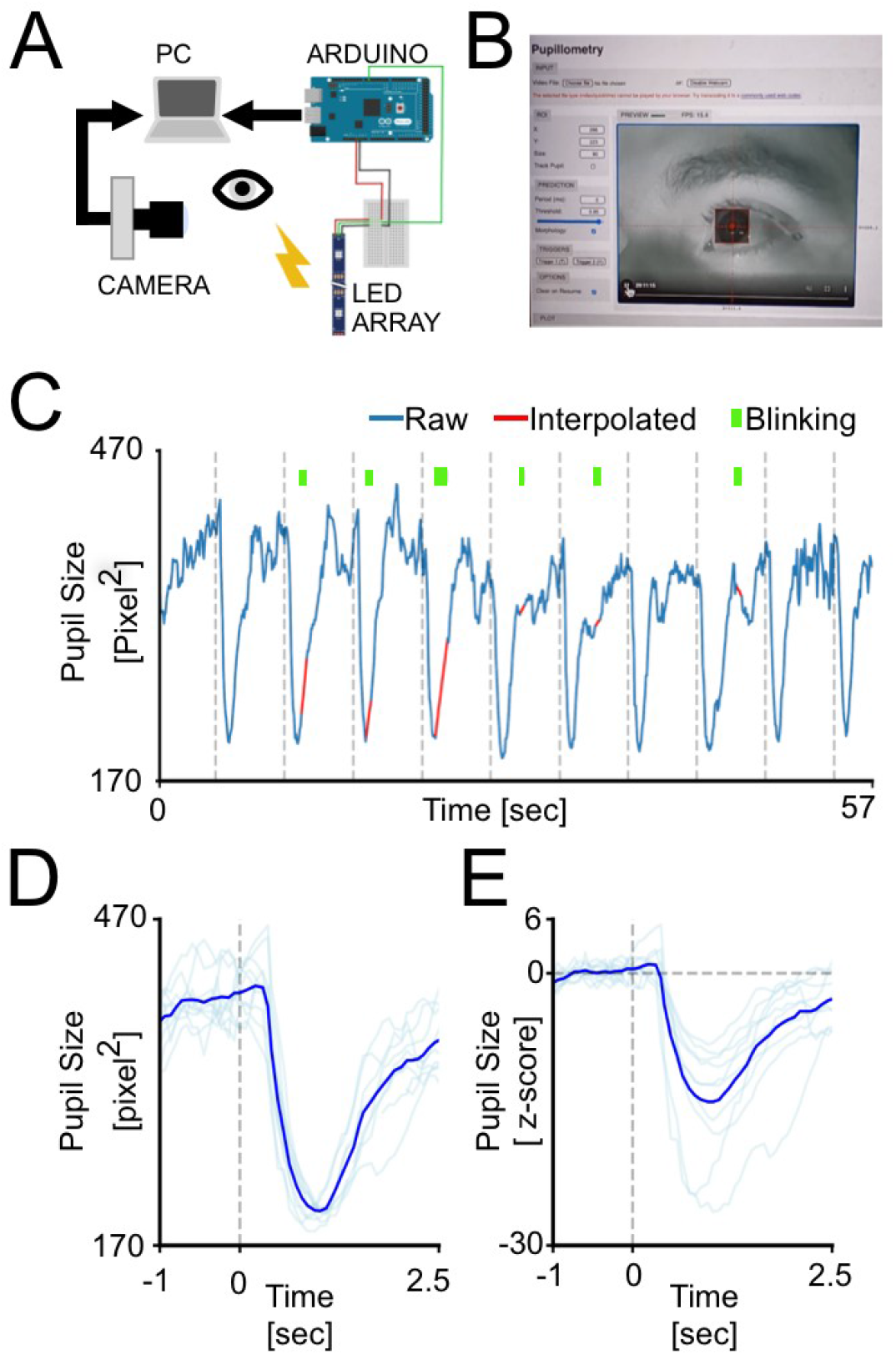
Web-browser Pupillometry Experiment. **A:** Experimental setup for running the PLR stimulation and in the meantime perform pupillometric recordings. The PC is connected to the internet running an instance of MEYE tool in the web browser. A USB camera, equipped with an IR light source, is focused on the eye of the subject. The photic stimulus is delivered using a LED array driven by an Arduino Due. The Arduino is connected to the PC emulating a keyboard and sending keystrokes stimulus triggers to the MEYE tool. **B:** A picture of MEYE graphical user interface. The subject during the recording is visualized as a streaming video. A ROI is used to locate the eye and a preview of the estimation of the pupil is superimposed to the image of the subject. The GUI allows to set different parameters of post-processing (map thresholding and refinement via mathematical morphology). C: Raw trace of the experiment (blue). Dashed lines locate the onset of flash stimuli. The green rectangles locate the onset and duration of blinks. The samples corresponding to blinks are removed and linearly interpolated (in red). **D:** Average event related transient to flash stimulation in raw values. After the onset of the stimulus (dashed line) a strong constriction of the pupil is observed (44.53%). **E:** Z-score of the average event related transient seen in D. The average nadir amplitude is 14.59 standard deviations from baseline.

#### Oddball Paradigm Corecordings

To compare the performances shown by the CNN system with that of a state of the art commercial software, we coregistered pupillometry using MEYE and an EyeLink 1000 system while 9 (3 males, 6 females, average age 28.78 years) participants executed an oddball paradigm. The experiment was conducted in a quiet, dark room. The participant sat in front of a monitor screen (88×50 cm) at a distance of 100 cm, with their head stabilized by a chin rest. Viewing was binocular. Stimuli were generated with the PsychoPhysics Toolbox routines [41,42] for MATLAB (MATLAB r2010a, The MathWorks) and presented on a gamma-calibrated PROPixx DLP LED projector (VPixx Technologies Inc., Saint-Bruno-de-Montarville, Canada) with a resolution of 1920×1080 pixels, and a refresh rate of 120 Hz. Pupil diameter was monitored at 1kHz with an EyeLink 1000 system (SR Research) with an infrared camera mounted below the screen and recording from the right eye. The participant was instructed to maintain fixation on a small dot (0.5 deg) presented in the center of the screen for the whole duration of the recording (300 seconds). In this study, the visual stimuli consisted in the appearance of a high probability stimulus (80% of times) defined as “Standard” and a lower probability stimulus (20% of times) defined as “Target”. The Standard stimulus consisted in a 100% contrast-modulated annular grating (mean luminance 25 cd/m^2), horizontally orientated, spatial frequency of 0.5 cpd, with an inner and outer diameter of 1.5 and 5 deg, respectively. The edges of the annulus were smoothed by convolving the stimulus with a gaussian mask (sigma = 0.5 deg). The Target stimulus has the same parameters of the Standard stimulus except the orientation that was 45 deg (see Fig. 4 A). The presentation duration of each trial, either the Standard (0 deg) or Target (45 deg) trial, was 200 ms with the intertrial interval between two consecutive trials being 2800 ms. The phase of both the *Target* and the *Standard* stimuli was randomized across trials.The participants were instructed to press a button for a Target stimulus and not to respond for a Standard stimulus.

**Fig. 4:**
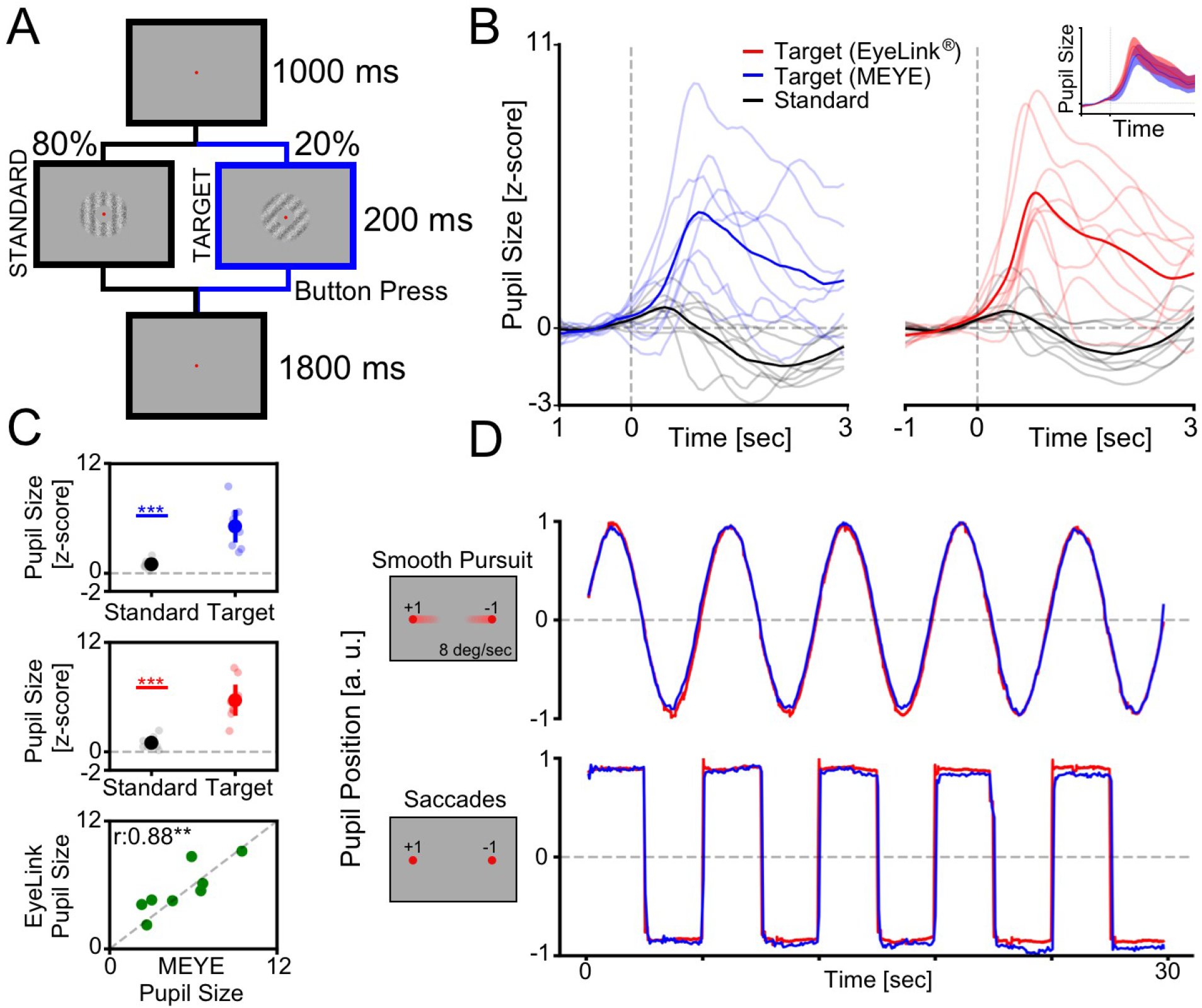
Cognitively driven pupillary changes. **A:** Visual Oddball procedure. The participant is instructed to fixate a small red dot placed at the center of the screen and to push a button only when the Target visual stimulus appears. **B:** Average Pupil waveforms. Average pupil response to Standard and Target stimulus for both MEYE (blue, left) and EyeLink (red, right). In the inset is represented the comparison between the evoked response to the Target stimulus in both setups. **C:** Average pupil response. Difference between the Standard and Target stimuli recording using MEYE (uppermost) and Eyelink (middle). The lowermost graph represents the correlation between MEYE and Eyelink data. **D:** Eye movements data. comparison between the MEYE tool(blue) and Eyelink system (red) during smooth pursuit task (upper) and saccades (lower).

**Fig. 5:**
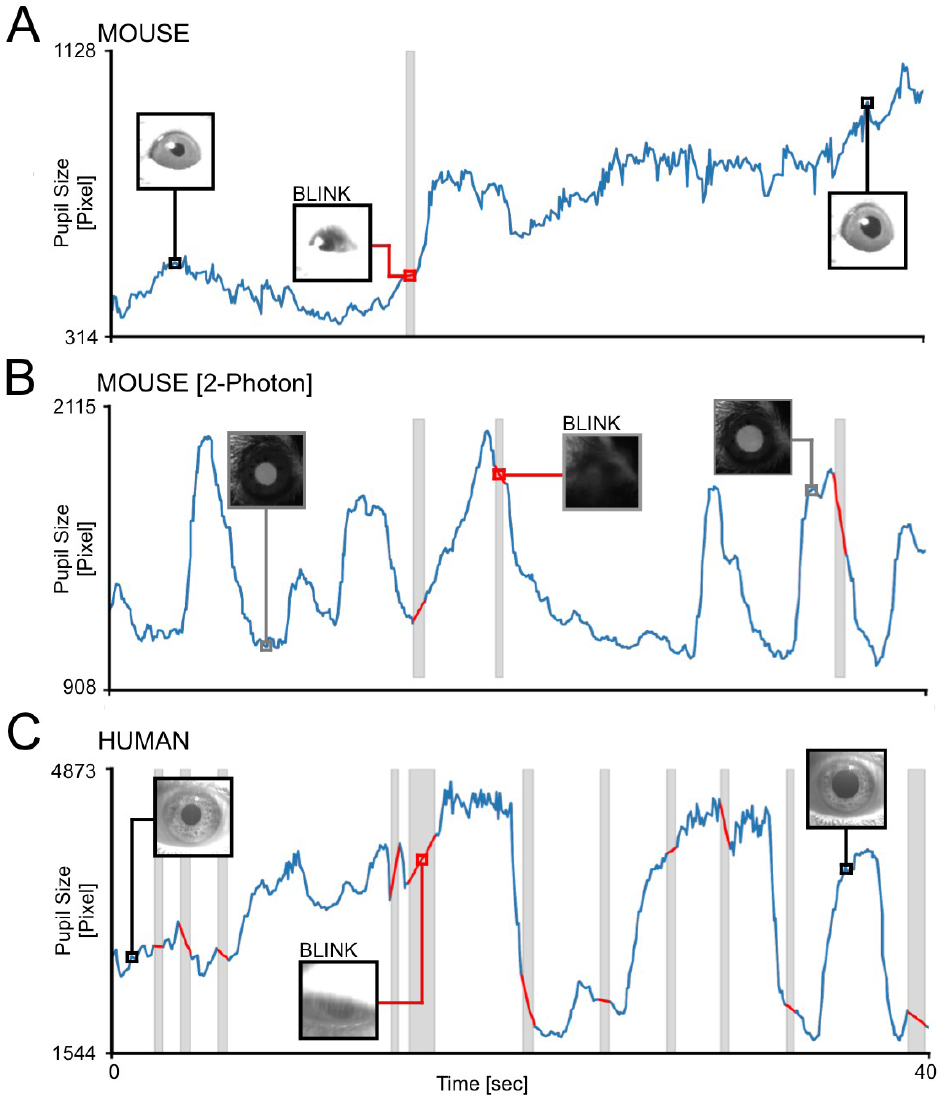
Offline Movies Analysis. **A:** Awake Head-fixed mouse running on a treadmill, recorded for 40 seconds. The grey area represents a blink, the trace of the blink is removed and linearly interpolated (Red line). **B:** Awake mouse during 2-photon calcium imaging. Here is clearly visible a brighter pupil with respect to A. Blinking epochs are removed and linearly interpolated. **C:** Pupillometry performed on a human subject, with a higher blinking rate with respect to mice. In all figures the insets images represent the ROIs.

#### Eye Movements Corecordings

For eye tracking recording we employed both the MEYE tool and EyeLink 1000 as described above. In the smooth pursuit condition a small dot (0.5 deg), moved on the screen horizontally, changing direction each 20 degrees of the visual field with a constant velocity of 8 deg/sec. In the Saccades condition every 2.5 s the small dot changes abruptly position horizontally with a span of 20 degrees.

## Data Availability

The code and web app are freely available on Github: github.com/fabiocarrara/meye

MEYE is available at: www.pupillometry.it

The dataset is available on: https://doi.org/10.5281/zenodo.4488164

## Results

### Pupillometry in Head-Fixed Mice

Our primary goal in this work is to examine if our CNN-based pupillometry can detect pupil fluctuations in real-time. We designed a behavioral experiment to characterize spontaneous and evoked pupillary changes. To confirm that pupil size changes are coupled with arousal transitions, we simultaneously recorded animal running speed during head fixation movements on the circular treadmill (Fig. 2 A). The experimental protocol included an initial period of habituation lasting 5 minutes followed by auditory stimulation using two tones (Tone1: 3 kHz, Tone2: 4kHz, using 120 seconds interstimulus, Fig. 2 B). We first set out to evaluate if CNN can detect event-related transients (ERT) due to auditory stimulation. For all the acoustic events, we averaged pupil size and running velocity during a 80 sec temporal window centered on acoustic stimulation (Fig. 2 C). We detected a significant pupillary dilation after the onset of the auditory stimulus, together with a similar peak in locomotion (Pupil: P-value < 0.01, Permutation Test, Velocity: P-value < 0.01, Permutation Test). This event-related transient is a proxy of the arousal change due to acoustic detection and can be considered a manifestation of cognitive and emotional processing of the stimulus [2]. By analyzing the overall traces, we found that the pupil diameter and the trace related to animal running, were characterized by a significant correlation (r: 0.79, P-value < 0.001, Pearson Correlation), in agreement with the hypothesis that fluctuations of the arousal level mainly characterize spontaneous pupillary events. Moreover, we calculated pupil size during habituation in different arousal states, finding that during locomotion, pupil size is significantly larger than in stationary periods (P-value < 0.01, Paired T-Test, Fig. 2 D). These results indicate that CNN pupillometry can detect spontaneous and elicited pupillary changes and can be used to monitor the mouse’s arousal state during head fixation experiments.

### Web-Browser Application to Perform Pupillometry Experiments

To test the implementation of the CNN in a web-browser (MEYE, Fig. 3 B), we designed a simple experiment aimed to measure PLR evoked by brief flashes of light on the human eye. The experiment included 10 flash events with an interstimulus of 5 seconds (dashed vertical lines in Fig. 3 C). The results showed a clear light-induced modulation of pupil size in correspondence with each flashes onset. Aligning and averaging all the traces along with the events, PLR can be quantified in both the raw (44.53%±0.67% change from baseline) and z-scored (14.59±2.05 st.dev. from baseline) trace (Fig. 3 D-E). To detect if it is possible to measure cognitively driven pupil signals using the MEYE tool reliably, we performed pupillometry while participants executed an oddball task, a commonly used paradigm for cognitive and attentional measurement. This task is based on the principle by which pupil dilation is stronger in response to rare stimuli and can be used as a physiological marker for the detection of deviant stimuli [37]. This experiment has been carried out recording the same eye using both the MEYE tool and an EyeLink 1000 system. According to Google Scholar, the Eyelink system is one of the most utilized eye trackers in psychology, psychophysics, and neuroscience, with more than 17K scientific publications mentioning this tool. During the oddball experiment, the subject was instructed to maintain fixation on a small dot presented in the center of the screen, pushing a button only when the *Target* stimulus appears on the screen and not responding to the *Standard* stimulus (Fig. 4 A). Averaging and comparing the responses to *Standard* and *Target* gratings results in a significant stronger pupil dilation for the *Target* stimulus than the *Standard* stimulus, that is detected by both the recording systems (MEYE: P-value < 0.001, T-Test Paired, EyeLink: P-value < 0.001, T-Test Paired, Fig. 4 B-C). No differences have been found for the responses evoked by the *Target* stimulus between the MEYE tool and the EyeLink system (P-value:0.327, T-test Paired, Fig. 4 B-inset). Moreover, the single-subject pupillary evoked amplitudes show a significant positive correlation between the two techniques (P-value:0.01, r:0.88, Pearson Correlation) with more than 75% of the variability explained by the linear model. Pupil-size is known to covary with eye position in video-based measurements [43], producing foreshortening of the pupillary image because the camera is fixed but the eye rotates. To overcome this issue, there are several possible solutions: the most simple requires to maintain constant fixation throughout each trial, but, if this requirement cannot be satisfied (such as in sentence reading), the position of the pupil at each sample is required to correct and mitigate the estimation error. Thus, we decided to quantify the agreement between positional outputs provided by MEYE and Eyelink for horizontal eye movements. We designed two tasks: in the first task, a dot smoothly traveled horizontally on the screen from left to right and vice versa at a velocity of 8 deg/sec and spanning 20 degrees, producing slow and smooth pursuit eye movements. In the other experiment, a dot jumped every 5 seconds from a position to the other (spanning 20 degrees), producing large, fast, and abrupt saccades. Results (Fig. 4 D) show that smooth pursuit movements generate a sinusoidal change of position with a good agreement between both systems (MAE: 0.04). The second task, inducing saccades, produces a slightly larger error (MAE: 0.073). This error is mainly due to the much lower sampling rate of MEYE (MEYE:15 fps; Eyelink: 1000 fps). This means that even if MEYE provides the exact positional information for each sample, it has a lower performance in adequately describing fast eye movements, such as saccades. Thus, MEYE provides the data required for post-hoc correction of pupil measures although it should be used with caution for measuring saccades. This factor should be taken into account when designing experiments using MEYE. Finally, MEYE can also be used as an offline tool to analyze pupillometry videos in various file formats, depending on the video codec installed in the web-browser. We successfully analyzed videos in different experimental conditions, including head-fixed mice running on a treadmill, during 2-photon calcium imaging, and in humans (Fig. 4 A-C).

## Discussion

In this work, we demonstrated that MEYE is a sensitive tool that can be used to study pupil dynamics in both humans and mice. Furthermore, by providing eye position MEYE allows post-hoc control of possible effects of eye movements on pupil measures[43]. MEYE can detect both spontaneous and evoked pupil changes in a variety of conditions: mice with black pupils in normal illumination conditions, and mice with bright pupils resulting from laser infrared illumination. This flexibility allows the use of MEYE in combination with 2-photon, wide field imaging and electrophysiological techniques widely adopted in the awake or anaesthetized mice. Furthermore, MEYE can be employed to design standalone experiments using cost-effective hardware with performance comparable with that of state-of-the-art commercial software. In this experiment, we used a USB webcam with a varifocal objective that allows focal adjustment concentrated on the eye. The cost of the imaging equipment is less than 50 euros (see Table 2), and requires no knowledge of coding to set up. The flashing stimulus apparatus requires a basic understanding of Arduino boards and can be assembled at a price lower than 50 euros. The overall cost of the apparatus is less than 100 euros. Our code can be used in two different ways, to satisfy many needs. One way relies on the standalone web-browser tool, that allows running MEYE on almost any device, from scientific workstations to notebooks or even smartphones. The other way utilizes a dedicated Python script running the CNN locally on a workstation. This latter case is suited for experiments with specific requirements, like high and stable framerate or online processing of pupil size in which on-the-fly pupil computer-interaction is required.

**Table 1.** Head-fixation apparatus components

| Part Number | Description | Qty | Price(euro) |
| --- | --- | --- | --- |
| TRB1/M | Locking ball and socket mount M4 | 1 | 55.83 |
| TR150/M-P5 | Optical post M4 - M6 150mm 5pack | 1 | 29.97 |
| RA90/M-P5 | Right-Angle Clamp | 1 | 45.7 |
| MB2020/M | Aluminium breadboard | 1 | 72.3 |
| RS075P/M | Pedestal Pillar Post | 1 | 21.63 |
| SS4MS12 | Setscrews 12mm long M4 | 1 | 5.61 |
| AP4M3M | Adaptor M4-M3 | 5 | 1.91 |
| ER025 | Cage assembly rod | 5 | 4.73 |
| SS6MS12 | Setscrews 12mm long M6 | 1 | 5.55 |
| CF038C-P5 | Clamping Fork | 1 | 46.49 |
| TOTAL |  |  | 289.72 |

**Table 2:** Hardware equipment for PLR.

| Part Number | Description | Qty | Price(euro) | store | Manufacturer |
| --- | --- | --- | --- | --- | --- |
| Walfront5k3psm<br>v97x | USB webcam | 1 | 33.48 | Amazon | Walfront |
| 149129 | Varifocal M12 Lens | 1 | 12.03 | Amazon | Sodial |
| A000062 | Microcontroller<br>Arduino Due | 1 | 35 | Arduino<br>Store | Arduino |
| 1312 | 4 NeoPixel RGB LEDs | 1 | 6.53 | Adafruit | Adafruit |
| Total Amount |  |  | 87.04 |  |  |

Valid open source and commercial alternatives exist, most of them are dedicated to gazing tracking and/or pupillometry. Commercial options are costly (tobii.com, sr-research.com, neuroptics.com), whereas open-source code instead requires programming knowledge and most of them are explicitly dedicated to one species [33,34]. One of these papers [33] assessed pupil dilation in mice through DeepLabCut [44], a technique for 3D markerless pose estimation based on transfer learning. This approach, albeit powerful, is conceptually different, since it is trained on user-defined key-points instead of using the entire pupil to perform semantic segmentation. The former technique is more suited to track and locate arbitrary objects on an image, the latter technique is focused on a more precise quantification of even small changes of the object area, since pixel-wise segmentation masks are refined iteratively using local and global context. The possible contribution of the web app technology resides in its portability: no software needs to be manually installed and configuration is minimal. Only a clear IR image of the subject’s eye is required. The performances of the tool are dependent on the host computer but it runs at >10 fps in most of the machines tested. This advantage is particularly useful for settings with limited resources and space or for educational purposes. Web browser embedded pupillometry will also be crucial for human scientific research, clinical and preventive medicine. It would also be a promising tool in the recently growing field of telemedicine given its minimal setup that can run on an average notebook or even on a smartphone, it allows possible large-scale recruitment of subjects directly in their own homes. This greatly facilitates infants, psychiatric, and motor-impaired patients’ compliance, particularly for longitudinal research designs. We also released an open-source database of eyes composed of more than 11.000 images in various settings: head-fixed mice (black pupil), head-fixed two-photon imaging mice (white pupil), and human eyes. This dataset will grow over time to introduce new species and new use cases to increase, update, and strengthen MEYE performances. The possible scenarios can be further expanded in the future, due to the dynamic nature of CNN. It can be updated from the source, providing instantaneous updates on each computer running an instance of the program. Our hope is to create a community that refines and consolidates pupillometric performances, to produce a tool that can be applied in different environments.

## Authors’ contributions

RM, FC and TP designed the research; RM, FC and GA trained and designed the AI tools; AV, RM, LL, GR and LLV labeled manually the dataset, RM, AV, LL, GS, LLV and AB performed the experiments, RM and FC performed data analysis.

## Acknowledgments

We gratefully acknowledge NVIDIA Corporation’s support with the Jetson AGX Xavier Developer Kit’s donation for this research. Authors would also like to thank Dr. Viviana Marchi, Dr. Grazia Rutigliano and Dr. Carlo Campagnoli for the critical reading of the manuscript.

